# Exploring Comorbidity Networks in Mild Traumatic Brain Injury Subjects through Graph Theory: A Traumatic Brain Injury Model Systems Study

**DOI:** 10.1101/2024.07.02.601618

**Authors:** Kaustav Mehta, Shyam Kumar Sudhakar

## Abstract

Traumatic brain injuries (TBIs) are characterized by myriad comorbidities that affect the functioning of the affected individuals. The comorbidities that TBI subjects experience span a wide range, ranging from psychiatric diseases to those that affect the various systems of the body. This is compounded by the fact that the problems that TBI subjects face could span over an extended period post-primary injury. Further, no drug exists to prevent the spread of secondary injuries after a primary impact. In this study, we employed graph theory to understand the patterns of comorbidities after mild TBIs. Upon application of network analysis and a novel clustering algorithm, we discovered interesting associations between comorbidities in young and old subjects with the condition. Specifically, bipolar disorder was seen as related to cardiovascular comorbidities, a pattern that was observed only in the young subjects. Similar associations between obsessive-compulsive disorder and rheumatoid arthritis were observed in young subjects. Psychiatric comorbidities exhibited differential associations with non-psychiatric comorbidities depending on the age of the cohort. The study results might have implications for effective surveillance and the management of comorbidities post mild TBIs.

## Introduction

Traumatic brain injuries (TBIs) are a significant public health concern affecting numerous countries across the globe (Gururaj 2002; Langlois et al. 2006; Maas et al. 2017; Rao et al. 2018; Dewan et al. 2019). TBIs, being one of the major causes of fatalities and disabilities, exerts a severe influence on the functioning of the affected population and their family members. Further, TBI survivors often have to deal with long-lasting complications that could significantly impact their quality of life and daily functioning (Brooks et al. 1986; Shoumitro et al. 1999; Hibbard et al. 2000; Maas et al. 2017; Sudhakar et al. 2023). The cost of diagnosing, treating TBIs, and managing post-traumatic complications in survivors of the condition could be enormously high, resulting in severe stress on the healthcare systems of the country (Langlois et al. 2006; Maas et al. 2017; Rao et al. 2018). Hence, a more significant focus and resources should be dedicated to enabling disease prevention.

TBI survivors often experience the development of a wide range of psychiatric and medical comorbidities that severely derail their work life and cognitive functioning (Brooks et al. 1986; Hibbard et al. 2000; Hammond et al. 2019; Sudhakar et al. 2023). These comorbidities are reported in TBIs of all severities, including mild, moderate, and severe forms of the injury (Hammond et al. 2019; Sudhakar et al. 2023). In a study of mild TBI (mTBI) subjects (Sudhakar et al. 2023), the authors quantified the prevalence of psychiatric and medical comorbidities at five years post-injury and as a function of the biological sex and age of the subjects. Mild TBI survivors are prone to developing highly diverse comorbidities ranging from cardiovascular, neurologic, and respiratory to psychiatric. Similar results have also been reported after severe and moderate impacts to the brain (Hammond et al. 2019).

The complications and comorbidities that TBI survivors develop after an impact on the brain can be attributed to primary and secondary brain injuries (Werner and Engelhard 2007; Maas et al. 2017). While primary injuries often happen immediately after an impact on the brain, the timescale of secondary brain injuries can vary from hours after the impact to days, months, and potentially years after the primary impact (Werner and Engelhard 2007; Algattas and Huang 2013; Maas et al. 2017). The primary injuries that damage local neurons, glia, and blood vessels are often irreversible (Werner and Engelhard 2007; Algattas and Huang 2013; Maas et al. 2017). On the other hand, the pathophysiological mechanisms guiding the development and subsequent spread of secondary brain injuries could be halted by applying numerous therapeutic strategies (Werner and Engelhard 2007; Algattas and Huang 2013; Prins et al. 2013). Unfortunately, despite the concerted efforts by the TBI community in search of a potential therapeutic agent to prevent the cascade of secondary brain injuries, no clinical trials have been successful so far on humans (Bullock et al. 1999; Xiong et al. 2009; Stein 2015; Sudhakar et al. 2019; Sudhakar 2023). Even though numerous drugs have shown superior efficacy in preventing secondary injuries in animal models of TBIs, such an effect was not seen in human clinical trials (Bullock et al. 1999; Stein 2015).

When successful treatment options don’t exist to treat the condition (Bullock et al. 1999; Xiong et al. 2009; Stein 2015), the focus should be on the careful management of comorbidities for effective patient care (Vella et al. 2017; Maas et al. 2017; Dash and Chavali 2018). However, continued surveillance of TBI survivors for all possible sets of comorbidities is impractical due to the associated costs and the wide range of comorbidities that TBI survivors often develop over a very long period following a primary impact (Brooks et al. 1986; Hibbard et al. 2000; Maas et al. 2017; Hammond et al. 2019; Sudhakar et al. 2023). The heterogeneous nature of TBI injury types also complicates the situation (Saatman et al. 2008; Maas 2016). TBIs are considered very heterogeneous owing to several factors, and it is widely expected that heterogeneous injuries could give rise to diverse patterns in the development of comorbidities over time (Saatman et al. 2008; Maas 2016). Hence, a more practical and economical way for surveillance of comorbidities in TBI subjects is required to prevent the occurrence of injury-associated complications (Sudhakar and Mehta 2024).

In this study, we propose to apply the principles of graph theory to decipher distinct patterns in the development of comorbidities after mild TBIs (Sudhakar and Mehta 2024). We analysed mTBI subjects five years post-injury using the Traumatic Brain Injury Model Systems National Database (TBIMS) data (Traumatic Brain Injury Model Systems Program 2021). By utilizing the data in the database (Traumatic Brain Injury Model Systems Program 2021), we constructed network graphs comprising comorbidities as nodes and edges representing the associations between them (Sudhakar and Mehta 2024). We decided to focus on mTBIs since the field is relatively under-explored, and the diagnosis of comorbidities post the condition goes largely unnoticed (Dean and Sterr 2013; Danna-Dos-Santos et al. 2018). Upon application of various statistical measures and clustering techniques, we uncover unique patterns in the development of post-traumatic comorbidities separately in young and old subjects. Our study yields novel observations in the development of comorbidities post mTBIs that could effectively be utilized for disease prevention and management.

## Methods

Our study employs graph theory approach (Sudhakar and Mehta 2024) to investigate patterns of disease co-occurrence following mTBIs (5 years). We leveraged the Traumatic Brain Injury Model Systems (TBIMS) database (Traumatic Brain Injury Model Systems Program 2021), a comprehensive database of TBI subjects encompassing longitudinal information about the development of comorbidities. Specifically, we included subjects with mTBIs as determined by the Glasgow Coma Scale (GCS) score (13<=GCS<=15) (Maas et al. 2017; Jain and Iverson LM 2022). We also restricted our analysis to patients with five years of follow-up data to capture a sufficient time frame for potential disease co-occurrences to emerge. A request to access the data in the public version of the database (Traumatic Brain Injury Model Systems Program 2021) was placed in February 2023, and the access was granted subsequently.

The information regarding possible comorbidities (presence, absence, and onset) was collected through the National Health and Nutritional Examination Survey (NHANES) (Center for Disease Control and National Center for Health Statistics 1999). A total of 26 medical and psychiatric comorbidities were included in our study, which were grouped under the following categories: psychiatric, musculoskeletal and rheumatologic, cardiovascular, respiratory, neurologic, endocrine, gastrointestinal, and ophthalmologic group of comorbidities. Under psychiatric comorbidities, the following comorbidities were included: Alcoholism, drug addiction (DA), depression, anxiety, obsessive-compulsive disorder (OCD), panic attacks (PA), bipolar disorder (BPD), attention deficit disorder/ attention deficit hyperactivity disorder (ADDADHD), and post-traumatic stress disorder (PTSD). The cardiovascular group of comorbidities had information regarding hypertension, congestive heart failure (CHF), stroke, myocardial infarction (MI), high blood cholesterol (HBC), and other heart conditions (OHC). Sleep disorder (SD), dementia, and movement disorder (MD) were included under the neurologic category. The musculoskeletal and rheumatologic groups include rheumatoid arthritis (RA), osteoarthritis (OA), and chronic pain (CP). Diabetes, cataracts, and liver disease (LD) were included under endocrine, ophthalmologic, and gastrointestinal conditions, respectively.

### Construction of the disease comorbidity network

Following the data filtering of mTBI subjects from the database (with age >=16 years), we computed phi correlation coefficient (φ) to quantify the strength of association between each unique pair of the 26 comorbidities (Fotouhi et al. 2018; Ljubic et al. 2020; Lee and Park 2021). φ is a non-parametric statistic suitable for analysing relationships between binary variables (Barabási et al. 2011; Fotouhi et al. 2018; Ljubic et al. 2020; Lee and Park 2021). To compute φ, we first constructed a 2*2 matrix for each pair of diseases (D1, D2) as per Table 1.

**Table 1:** Matrix for computation of phi correlation coefficients. a is the number of patients with both diseases D1 and D2, b is the number of patients with disease D1 but not disease D2, c is the number of patients with disease D2 but not disease D1, d is the number of patients with neither disease D1 nor disease D2. The values for N, e, f, g, and h are given by equation 1.

|  | **D1 = 1** | **D1 = 0** | **Total** |
| --- | --- | --- | --- |
| **D2 = 1** | a | c | g |
| **D2 = 0** | b | d | h |
|  | e | f | N |

In Table 1, a is the number of patients with both diseases D1 and D2, b is the number of patients with disease D1 but not disease D2, c is the number of patients with disease D2 but not disease D1, d is the number of patients with neither disease D1 nor disease D2. The values for N, e, f, g, and h are given by the following equation,

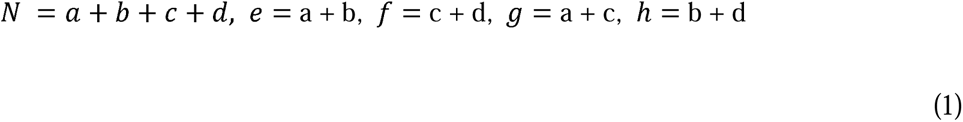

The phi coefficient (φ) is calculated using the following formula:

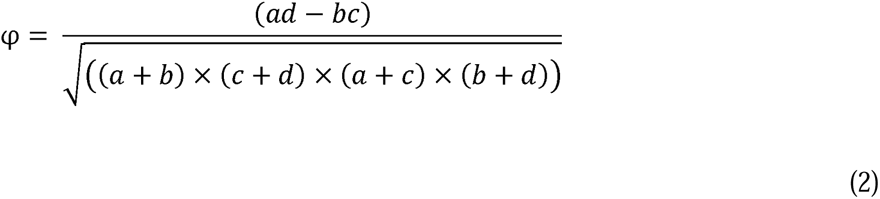

To assess the statistical significance of each phi correlation coefficient (Hidalgo et al. 2009), we performed a student’s t-test using the formula:

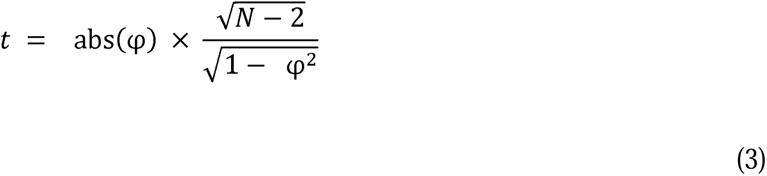

where N is given by equation 1. The calculated t-value for each disease pair was then compared against a critical t-value derived from an inverse t-distribution with N-2 degrees of freedom and a 99% confidence interval. For each pair of diseases, the critical t-values were generated for the upper tail for 99% probability and N-2 degrees of freedom. We, therefore, identified significant co-occurrences between comorbidities (magnitude) regardless of the direction of the relationship. If the calculated t-value exceeds the critical t-value, we reject the null hypothesis of no association and conclude that a statistically significant co-occurrence exists between the two diseases. This stringent significance level of 99% (p < 0.01) ensures that only robust co-occurrence patterns are included in the subsequent network analysis, minimizing the inclusion of spurious associations. See Figure S1 for a comprehensive map of significant and non-significant comorbidity associations.

Next, we constructed a disease comorbidity network (all subjects, young and old) using the NetworkX library (https://networkx.org/), with each of the 26 diseases represented as a node. Subjects with an age at follow-up (5 years) less than or equal to 50 years were categorized as young, and those over 50 years were classified as old. Edges were added between nodes that exhibited statistically significant co-occurrence based on the t-test results. The weight of each edge was assigned the absolute value of the corresponding Phi coefficient, reflecting the strength of the association (Fotouhi et al. 2018).

To further study the disease comorbidity networks (all subjects, young and old), several centrality measures (Klein 2010; Fotouhi et al. 2018; Bloch et al. 2021; Evans and Chen 2022) were calculated to identify critical nodes and potential hubs of co-occurrence. These measures are listed below:

*Degree centrality*: Degree centrality represents the number of edges connected to a node, reflecting the number of diseases a disease significantly co-occurs with (Klein 2010; Fotouhi et al. 2018; Bloch et al. 2021; Evans and Chen 2022). In a disease comorbidity network, a high degree of centrality indicates a disease that frequently co-occurs with many other diseases, potentially suggesting a central role in multimorbidity patterns. The degree centrality C(w) for node w □ V is defined as (Klein 2010; Fotouhi et al. 2018; Bloch et al. 2021):

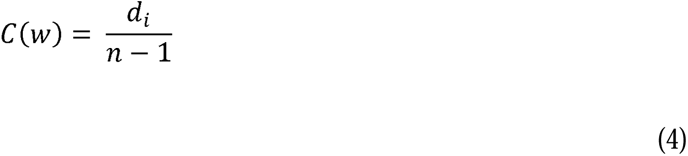

where *d_i_* is the total number of nodes connected to the node w and n is the total number of nodes in the graph V.

*Betweenness centrality*: Betweenness centrality represents the extent to which a node lies on the shortest paths between other nodes in the network (Klein 2010; Fotouhi et al. 2018; Bloch et al. 2021). Nodes that score high on betweenness centrality often act as a bridge or connector between different disease clusters, indicating their importance in the flow of information or disease progression within the network (Klein 2010; Fotouhi et al. 2018; Camarillo-Ramirez 2020; Bloch et al. 2021). Diseases with high betweenness centrality play a crucial role in understanding the underlying mechanisms of comorbidity and could be potential targets for therapeutic intervention in the TBI demographic (Fotouhi et al. 2018). For a graph G = (V, E), consider three distinct nodes u, v, w □ V. Then, let σ*_u,v_* be the number of shortest paths that connect u and v, and σ*_u,v_* (*w*) is the number of shortest u-v paths that go through w. The betweenness centrality C(w) for node w □ V is defined as (Klein 2010; Fotouhi et al. 2018; Bloch et al. 2021):

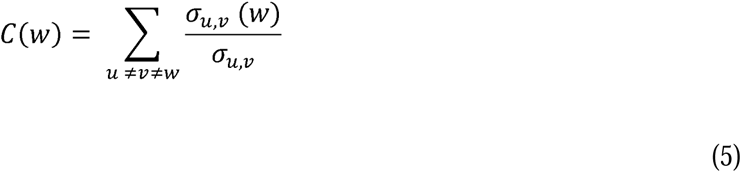

#### Eigenvector centrality

This measure considers the number of connections a node has and the centrality of its neighbours (Klein 2010; Fotouhi et al. 2018; Bloch et al. 2021). A node with high eigenvector centrality is connected to many other well-connected nodes (Klein 2010; Fotouhi et al. 2018; Bloch et al. 2021). In the context of disease comorbidity, a high eigenvector centrality suggests a disease that is frequently co-occurring and linked to other diseases with many co-occurrences, potentially indicating a pivotal role in complex disease networks. The eigenvector centrality for each node is calculated using the following formula (Klein 2010; Fotouhi et al. 2018; Bloch et al. 2021),

For a graph G = (V, E) with adjacency matrix A = *a_v,t_*, the eigenvector centrality score, *x_v_*, of a vertex v with its set of neighbours π(v) can be defined as:

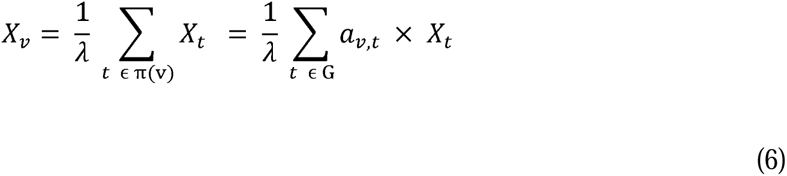

### BC Clustering Algorithm

To simplify the analysis and identify clusters of diseases that frequently co-occur in our network, we employed a betweenness centrality-based clustering approach adapted from Camarillo-Ramirez et al. (2020) (Camarillo-Ramirez 2020). This procedure was applied to the comorbidity networks of all three groups: all subjects, young and old subjects. We leveraged the NetworkX library (https://networkx.org/) to calculate betweenness centrality for each disease node in the network. The clustering algorithm iteratively removes nodes with the highest betweenness centrality values, acting as bridges between clusters. Removing nodes with high-betweenness centrality effectively splits the network into clusters of tightly connected modules that are less dependent on intermediary nodes for communication (Camarillo-Ramirez 2020). This process is continued until a predefined betweenness centrality threshold was reached, indicating the separation of distinct clusters (Camarillo-Ramirez 2020). We explored multiple thresholding strategies, including using the minimum, median, or user-specified values for flexibility, but chose to use the median betweenness centrality of the initial network.

Following identifying connected components within the remaining network, we employed a strategy to refine the clusters by considering previously removed nodes with high betweenness centrality (Camarillo-Ramirez 2020). We reintroduced such nodes back into the cluster if they maintained direct connections (edges) to at least one node within the identified connected component (Camarillo-Ramirez 2020). This step helped us capture potentially important diseases that could bridge clusters and influence the flow of information or co-occurrence of diseases within the network (Camarillo-Ramirez 2020). By incorporating these bridging nodes, the resulting clusters likely provide a more comprehensive and biologically relevant representation of the underlying disease relationships within our TBI-associated disease network (Camarillo-Ramirez 2020).

## Results

We analysed the information present in the TBIMS national database (Traumatic Brain Injury Model Systems Program 2021) to construct a disease comorbidity network of mTBI subjects at 5 years post injury. The total number of mTBI subjects at 5 years post injury is 4915. Altogether there was information regarding 26 comorbidities, as mentioned in the Methods section (Figure 1), collected through the NHANES survey (Center for Disease Control and National Center for Health Statistics 1999) the prevalence of which can be seen in a previous study (Sudhakar et al. 2023).

**Figure 1:**
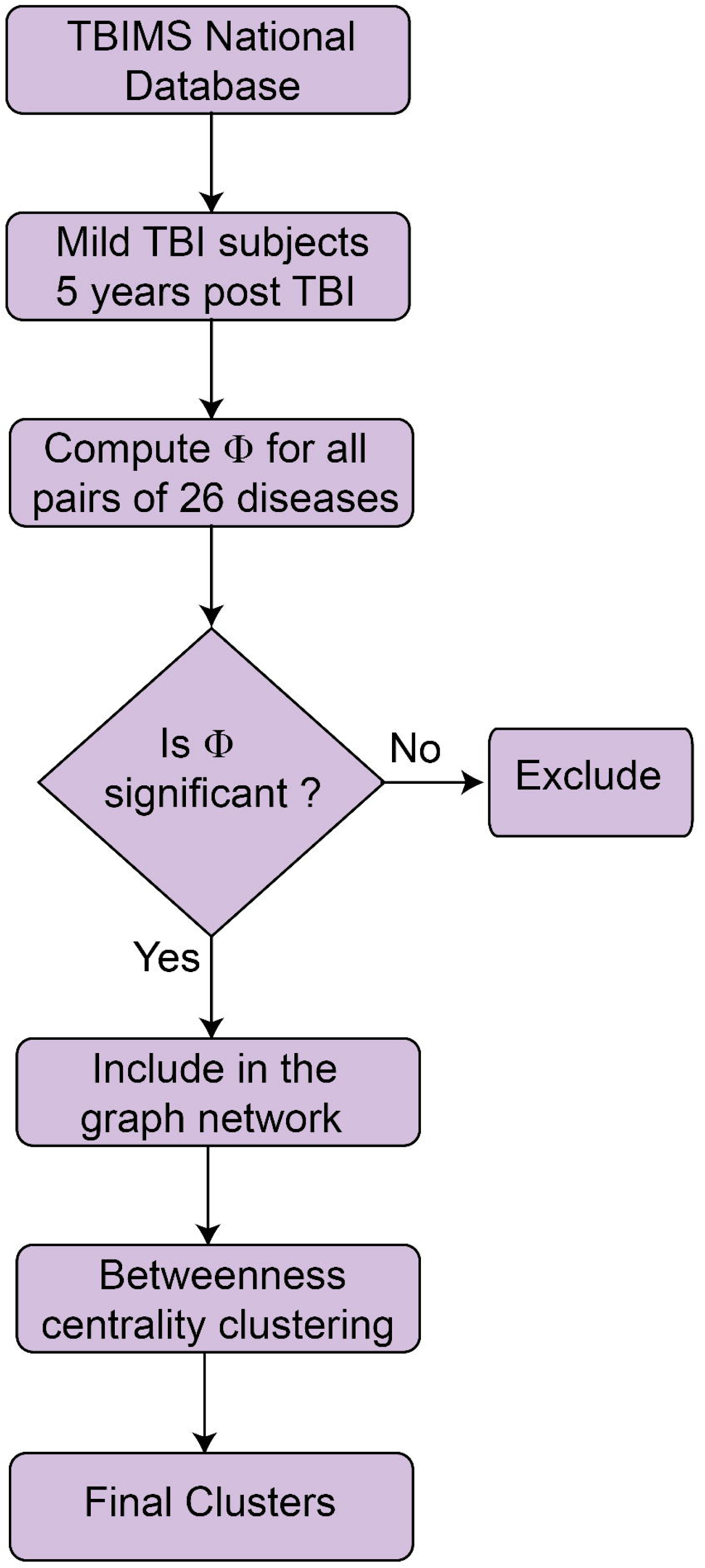
Schematic workflow of the analysis of disease comorbidity networks in mTBI subjects Flow chart represents various steps in the construction of disease comorbidity network and subsequent clustering based on betweenness centrality

As a first step, we computed the extent of co-association between the 26 medical and psychiatric comorbidities. To do so, we calculated the phi-correlation coefficient (φ) between each pair of comorbidities (Barabási et al. 2011; Fotouhi et al. 2018). After performing a statistical test, we eliminated weaker connections and retained only the statistically significant ones (Figure 1). The resulting comorbidities and their connections were represented in the form of a graph network where nodes represent the individual comorbidities (node size represents prevalence) and edges represent the associations in the form of computed φ correlations (Figure 2). Thicker edges represent stronger connections between comorbidities (Figure 2). For example, depression exhibits a stronger association with anxiety (φ = 0.48), one of the strongest associations between all sets of comorbidities in the network. On the other hand, chronic pain exhibits a somewhat weaker yet statistically significant connection with anxiety (φ = 0.21). Any isolated comorbidity (stroke) with no connection to other parts of the network was not included (Figure 2).

**Figure 2:**
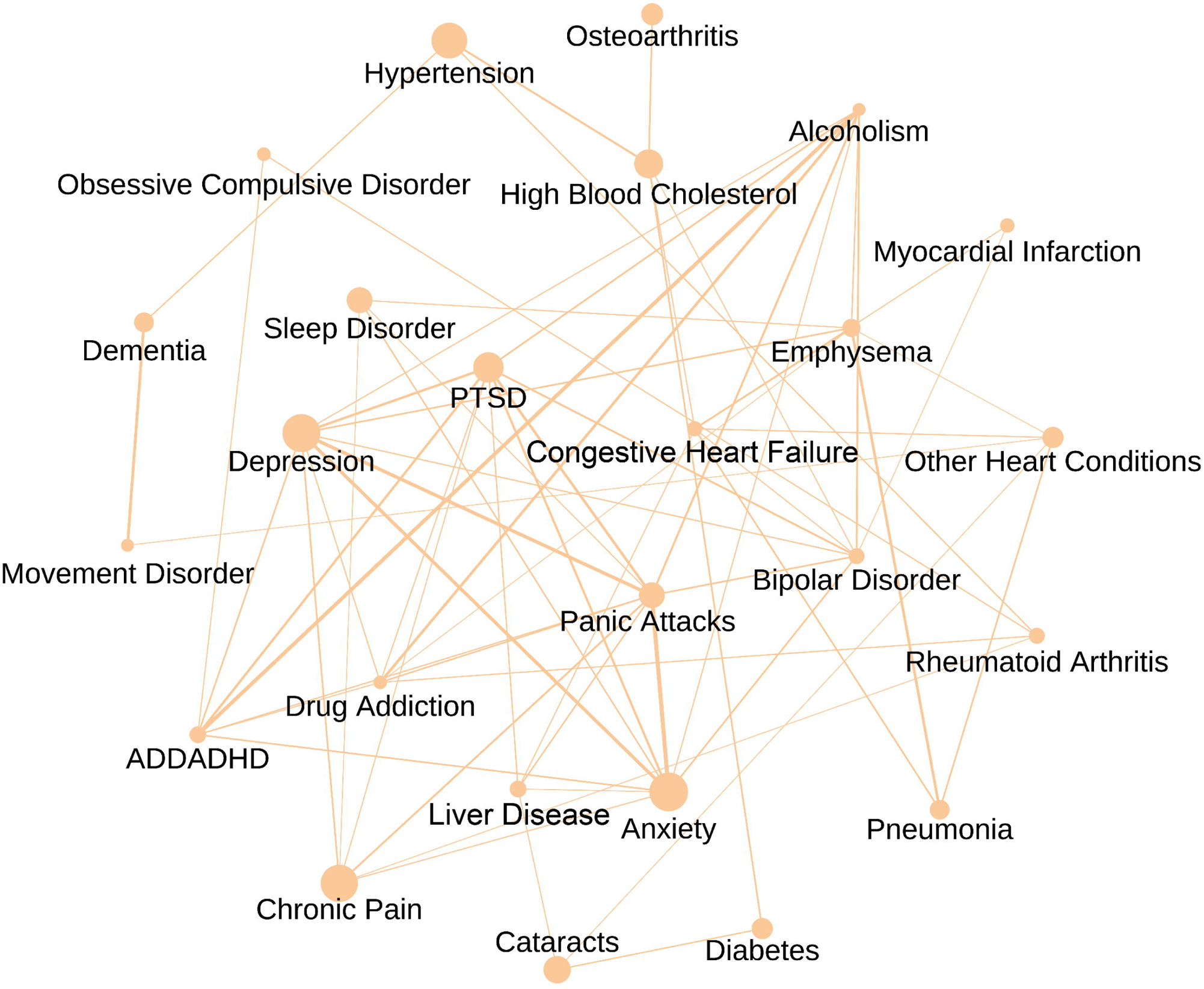
Disease comorbidity network of all mTBI subjects at 5 years post injury The nodes in the network represent the comorbidities while the edges represent associations in the form of phi-correlation coefficient. Node size indicates prevalence and edge thickness represent the strength of the association

To understand the statistical properties of the network and identify vital nodes (comorbidities), we computed various centrality measures (Klein 2010; Fotouhi et al. 2018; Bloch et al. 2021). Three necessary centrality measures were considered: betweenness centrality, degree centrality, and eigenvector centrality (see METHODS). All centrality measures identify vital nodes that control information flow and transition patterns in the disease comorbidity network (Fotouhi et al. 2018). The distribution of the three centrality measures of the network can be seen in Figure 3. From the figure, it can be seen that the three centrality distributions differ from each other, although a high correlation can be seen between degree vs eigenvector centralities (r = 0.93). Since betweenness centrality exhibited the most minor correlation with prevalence (r = 0.028), we investigated the network further using this metric.

**Figure 3:**
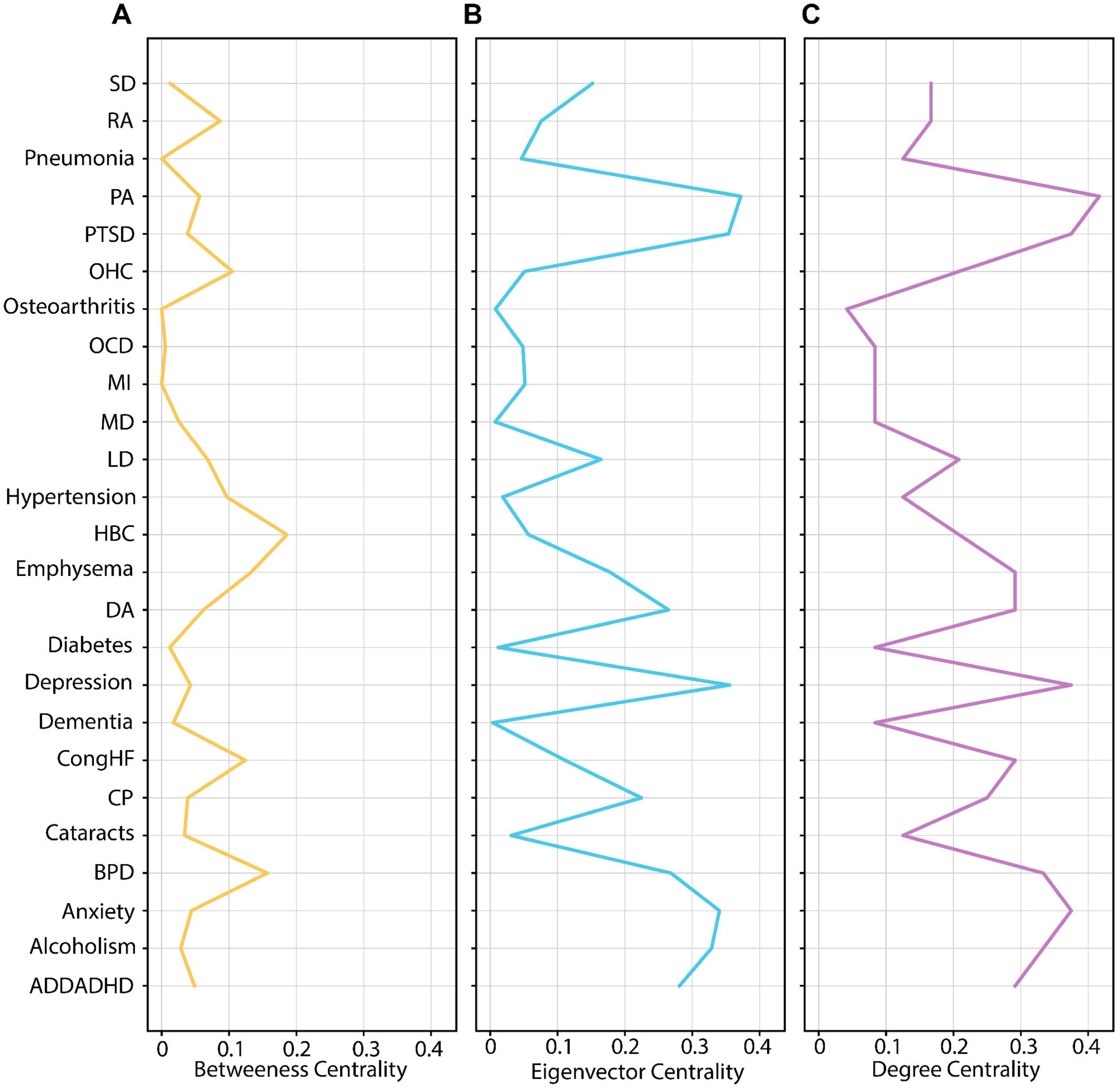
Distribution of three centrality measures constructed from the comorbidity network of all mTBI subjects. Panel A represents the distribution of betweenness centrality, panel B represents degree centrality and panel C represent eigenvector centrality

Next, we decided to decipher the network at a more granular level by identifying localized clusters of comorbidities after eliminating connections between vital comorbidity hubs. To do so, we incorporated an algorithm implemented in a recent study of disease comorbidities (Camarillo-Ramirez 2020). The algorithm identifies nodes with high betweenness centrality values and removes them iteratively from the network until a specific threshold is reached (Camarillo-Ramirez 2020). The algorithm then reinstates the link between the removed nodes and the remaining connected components in the graph, forming localized clusters of comorbidities (see Table 2 for removed nodes) (Camarillo-Ramirez 2020). This approach aims to identify comorbidities that occur in close association with each other, which would further help decipher their shared physiological mechanisms.

**Table 2:**
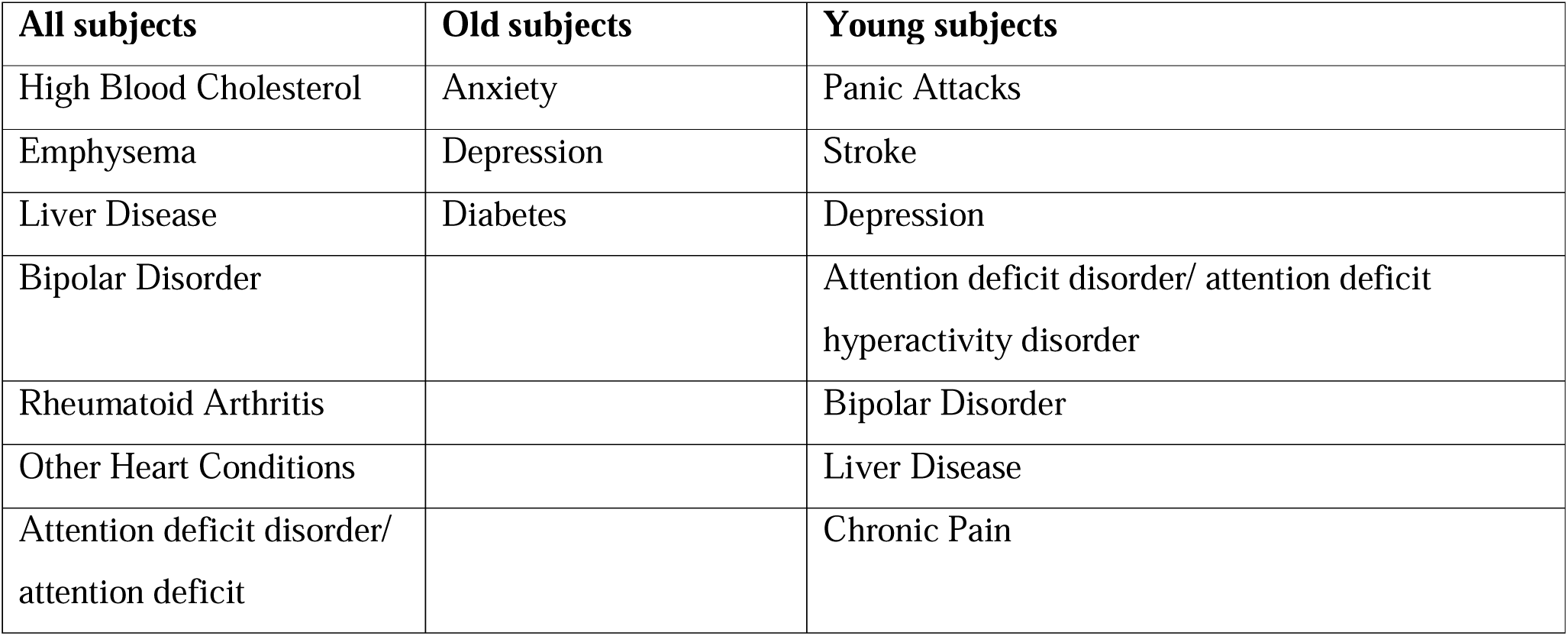
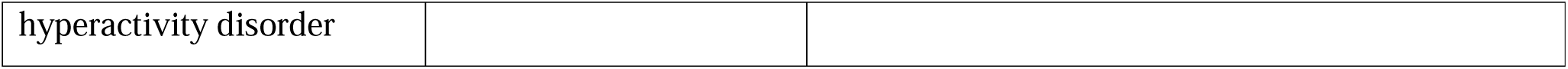
List of removed nodes applying the betweenness centrality algorithm to the comorbidity networks of all, young and old mTBI subjects.

### Comorbidity clusters in mTBI subjects

We identified six clusters (Figure 4) after applying the clustering method based on betweenness centrality (Camarillo-Ramirez 2020). The six clusters differed concerning the constituent nodes and connectivity between them. The first cluster (Figure 4) includes psychiatric conditions and their connections. The psychiatric conditions found in cluster 1 include PTSD, BPD, ADDADHD, PA, DA, alcoholism, anxiety, and depression (Figure 4). In addition to observing connections among psychiatric conditions, cluster 1 also encompasses connections between non-psychiatric and psychiatric conditions. The non-psychiatric conditions seen in cluster 1 include LD, RA, CP, SD, and emphysema. Mainly, emphysema is associated with depression, alcoholism, and DA. One salient point is that OCD is not associated with other psychiatric conditions in the network (except ADDADHD). Altogether, the first cluster seems to reinforce the fact that psychiatric conditions exhibit a high degree of comorbidity (Shoumitro et al. 1999; Hibbard et al. 2000; Sudhakar et al. 2023).

**Figure 4:**
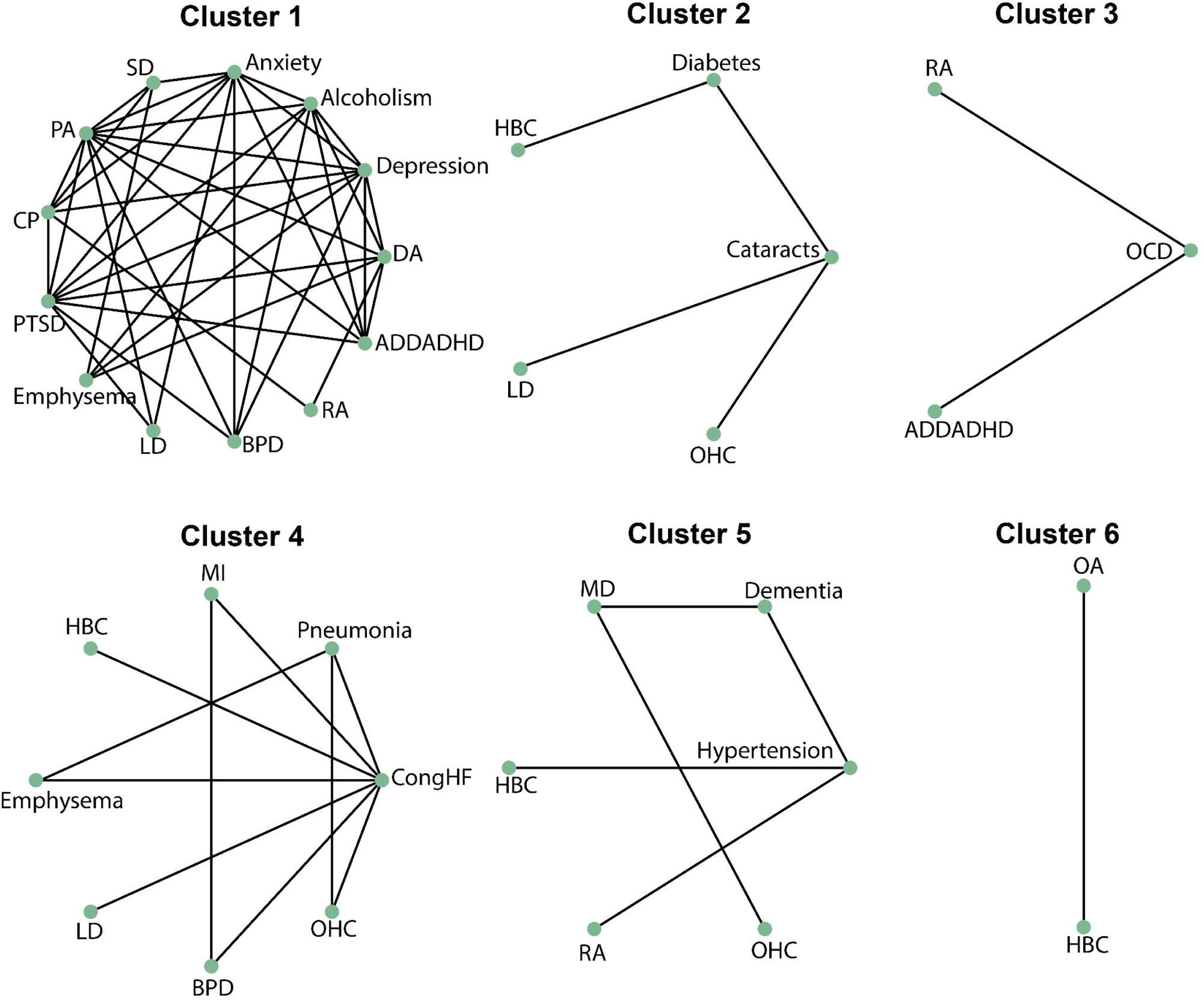
Application of betweenness centrality-based clustering to the comorbidity network of all mTBI subjects Individual clusters encompass nodes and edges that represent the comorbidity and the association between them respectively.

In the second cluster (Figure 4), diabetes and cataracts are associated with each other and other cardiovascular conditions. Diabetes is comorbid with HBC and cataracts with OHC. In addition, cataracts, an ophthalmologic condition, is associated with LD. The third cluster (Figure 4) consists of three constituents: OCD, ADDADHD, and RA. OCD, a psychiatric condition, is associated with both RA and ADDADHD. In the fourth cluster, one can witness the association between cardiovascular comorbidities and other groups of comorbidities. Cardiovascular comorbidities (CHF, OHC) in the cluster are associated with pulmonary comorbidities (pneumonia and emphysema). CHF is, in particular, comorbid with HBC, OHC, MI, LD, BPD, pneumonia, and emphysema. Lastly, BPD is associated with cardiovascular conditions CHF and MI.

In the cluster 5 (Figure 4), one can see the association between neurologic and cardiovascular conditions. In particular, dementia is associated with hypertension and MD is linked with OHC. Cluster 5 (Figure 4) also witnesses the association between MD and dementia, RA and HBC with hypertension. Lastly, an association between OA and HBC can be seen in cluster 6. In the next section, we will discuss which associations are seen in young and old mTBI subjects.

### Comorbidity clusters in old and young mTBI subjects

As a next step in the study, we wanted to ask which comorbidity patterns are observed in old and younger subjects. Subjects were categorized as old if their age was greater than 50 years at admission (or young otherwise). Similar to the analysis involving all mTBI subjects, we computed the φ for all pairs of 26 comorbidities of mTBI subjects (>50 years of age). After retaining the statistically significant associations, we displayed the information as a network graph (Figure 5) for old mTBI subjects. Note that stroke, MI, OA, and hypertension were excluded since they are not connected to the leading network. With this network in place (Figure 5), we applied the betweenness centrality clustering algorithm to identify localized clusters of comorbidities in the network.

**Figure 5:**
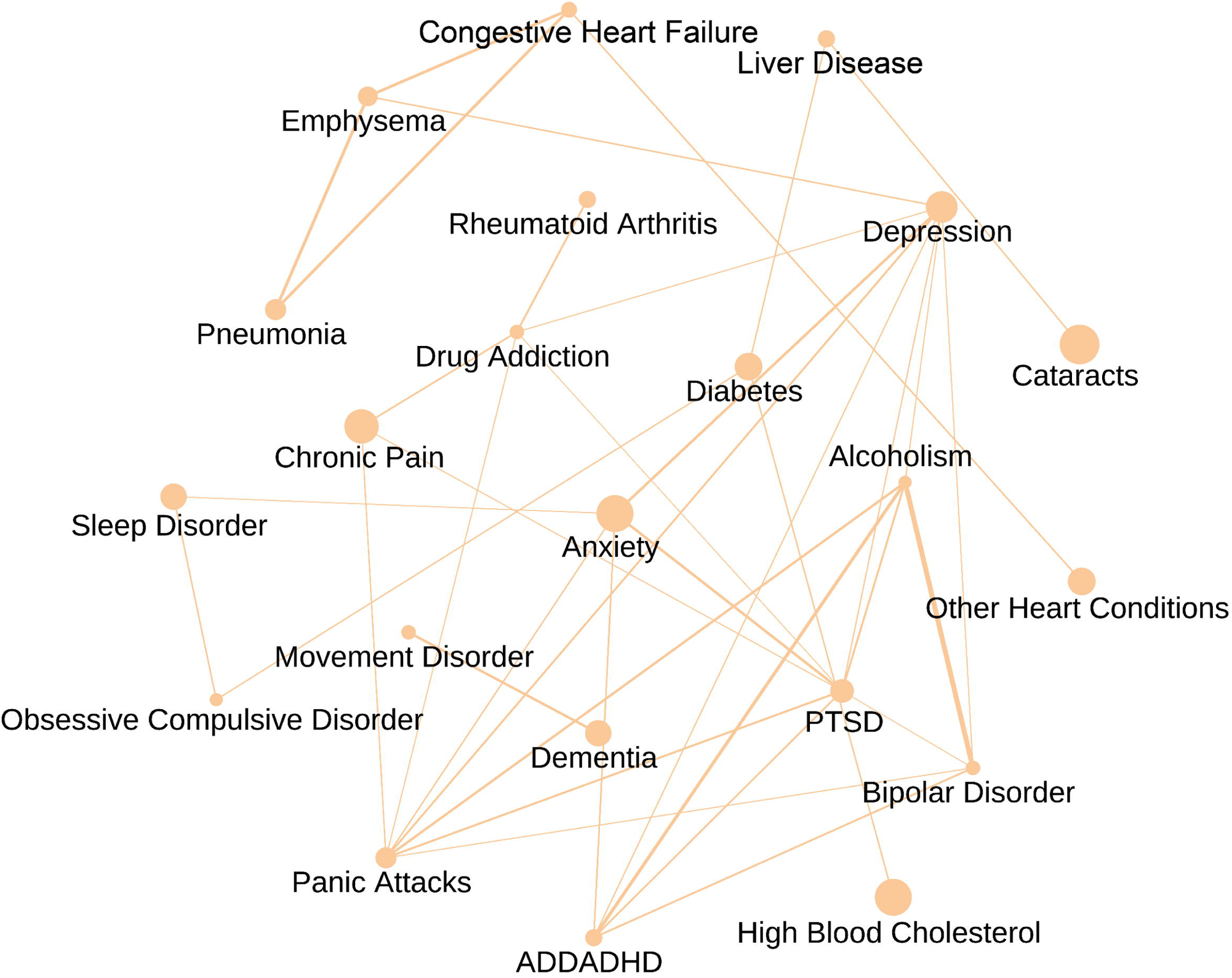
Disease comorbidity network of old mTBI subjects at 5 years post injury The nodes in the network represent the comorbidities while the edges represent associations in the form of phi-correlation coefficient. Node size indicates prevalence and edge thickness represent the strength of the association

First, we found that psychiatric conditions have a high degree of co-occurrence among each other similar to the results observed in the previous section. This can be seen from cluster 1 (Figure 6), where depression, anxiety, PTSD, DA, alcoholism, PA, and BPD tend to be co-associated with each other. In addition to co-occurrence within the group, psychiatric comorbidities are also associated with other somatic conditions. For example, CP is seen to be associated with DA, PA, and PTSD (cluster 1), while RA is associated with DA (cluster 1) and emphysema with depression (cluster 2, Figure 6). One can also witness co-association between CHF and pulmonary comorbidities (pneumonia and emphysema) in cluster 2 (Figure 6). Cluster 3 showcases the connection between dementia and MD, typically occurring as people age. Cluster 5 encompasses connections of OCD with diabetes and SD, while cluster 4 depicts the link between diabetes and HBC (Figure 6). Lastly, one can see that LD is associated with cataracts and diabetes in cluster 6. The list of removed nodes as a result of applying the clustering algorithm can be seen in Table 2.

**Figure 6:**
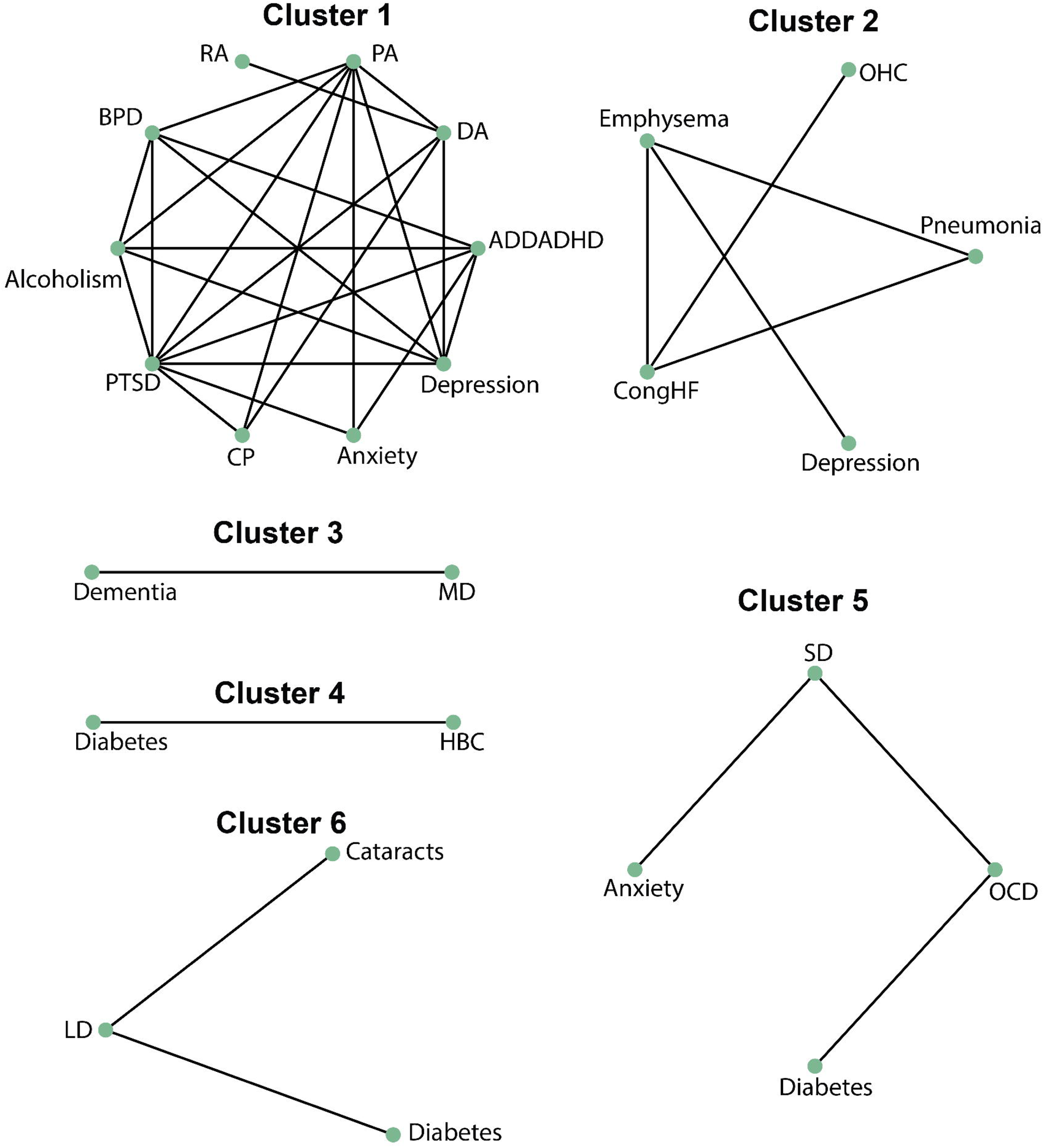
Application of betweenness centrality-based clustering to the comorbidity network of old mTBI subjects Individual clusters encompass nodes and edges that represent the comorbidity and the association between them respectively.

Next, we performed the same analysis for young mTBI subjects. After constructing the disease comorbidity network for this study group (Figure 7), we formed localized clusters by applying the clustering algorithm based on betweenness centrality as before. This resulted in seven clusters, as shown in Figure 8. Similar to the trends seen in older subjects, psychiatric comorbidities tend to co-occur (Figure 8). This can be seen across cluster 1 and in clusters 2 and 4 where anxiety and PTSD were connected to other psychiatric conditions respectively (Figure 8). Psychiatric comorbidities were also associated with other somatic diseases like emphysema (cluster 1), LD (clusters 2 and 4), and stroke (cluster 4) (Figure 8). We also observed a close association between BPD and cardiovascular comorbidities (cluster 3, MI, CHF, HBC). Cluster 5 (Figure 8) encompasses associations between OCD with RA and ADDADHD. Lastly, we observed that SD is associated with cardiovascular comorbidities (OHC, stroke) and CP (cluster 6).

**Figure 7:**
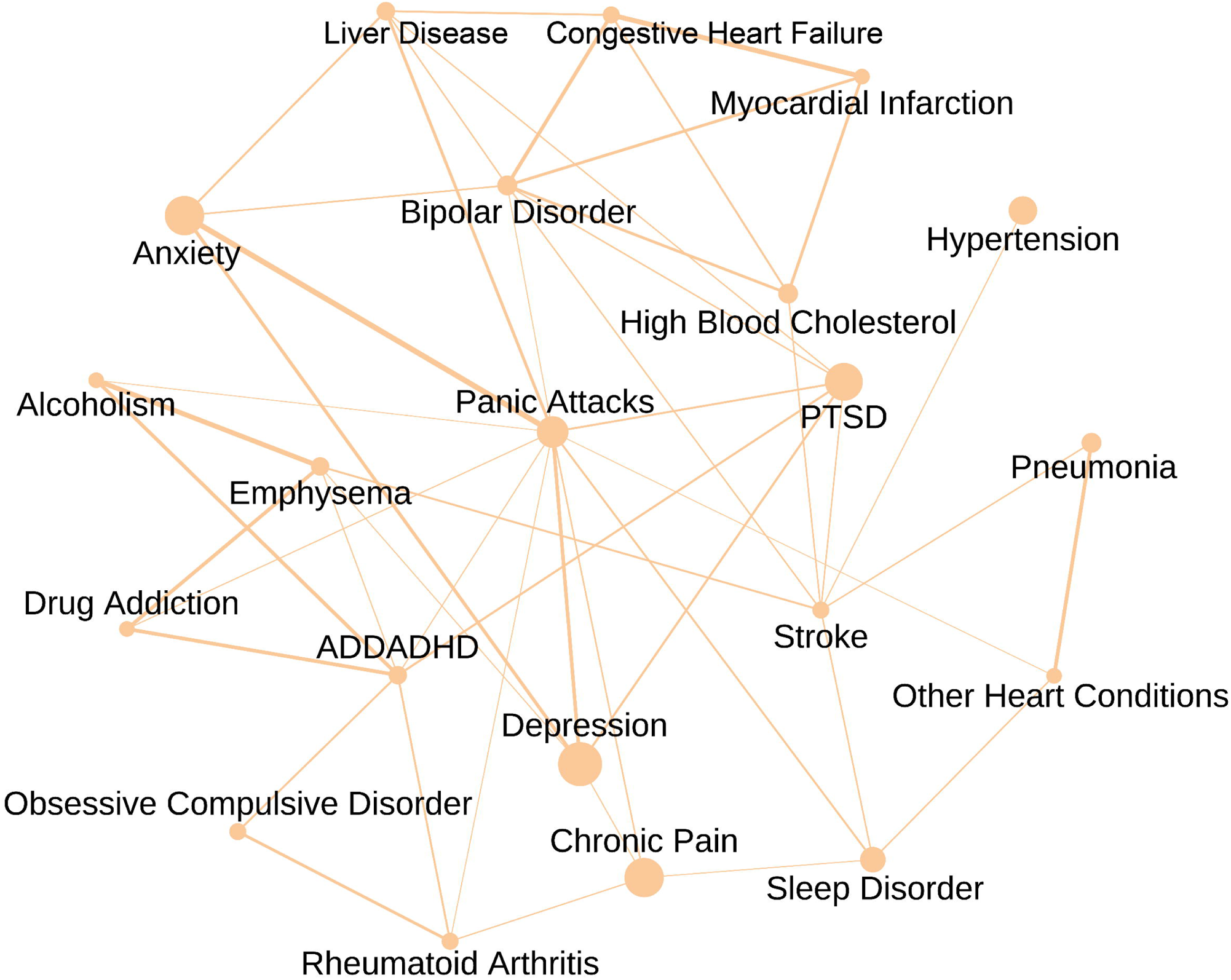
Disease comorbidity network of young mTBI subjects at 5 years post injury The nodes in the network represent the comorbidities while the edges represent associations in the form of phi-correlation coefficient. Node size indicates prevalence and edge thickness represent the strength of the association

**Figure 8:**
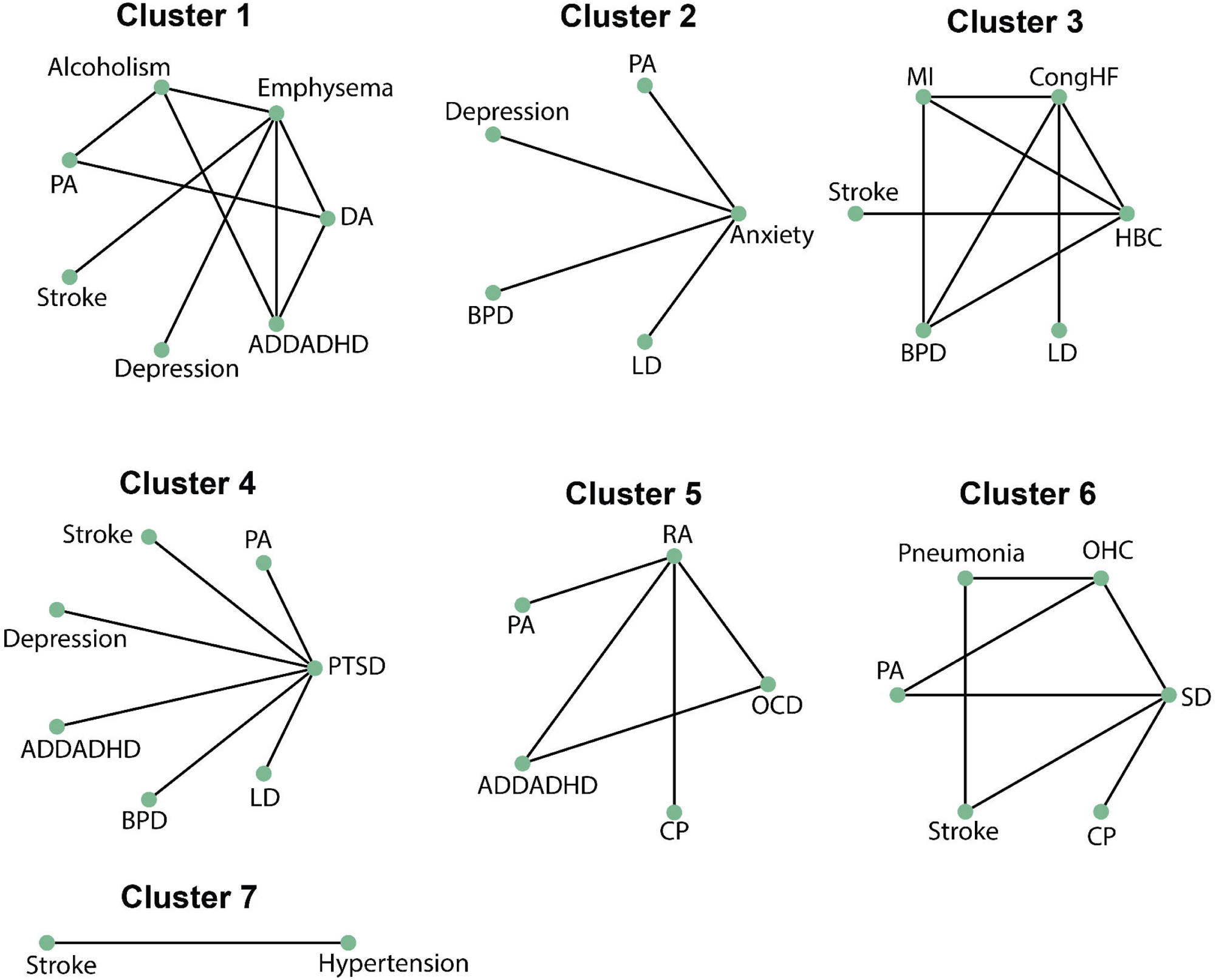
Application of betweenness centrality-based clustering to the comorbidity network of young mTBI subjects Individual clusters encompass nodes and edges that represent the comorbidity and the association between them respectively.

Next, we identified prominent comorbidity patterns in the network clusters of all subjects (Figure 2 and 4) and asked whether they were observed in the network of young and old subjects (Figure 6 and 8). The features are listed in Table 3, along with the information on whether they are observed in young vs old cohorts. The analysis revealed some striking similarities and dissimilarities regarding the co-occurrence of comorbidities between the two cohorts (Table 3). Both cohorts were characterized by solid co-occurrences of psychiatric comorbidities. However, the co-occurrence of psychiatric comorbidities with LD was seen only in the young cohort. The same co-occurrence with CP was witnessed only in the old cohort (Table 3). One remarkable feature of the disease comorbidity pattern in the young cohort is that OCD tends to co-occur with ADDADHD and RA (Table 3). In comparison, this association was not present in the old cohort. Cardiovascular comorbidities were associated with other groups of comorbidities in both young and old cohorts, but some differences were also noticed. The association of cardiovascular comorbidities with BPD was seen only in the young cohort (Table 3). The implications of these relative associations in young vs old cohorts are explained in the discussion section.

**Table 3:**
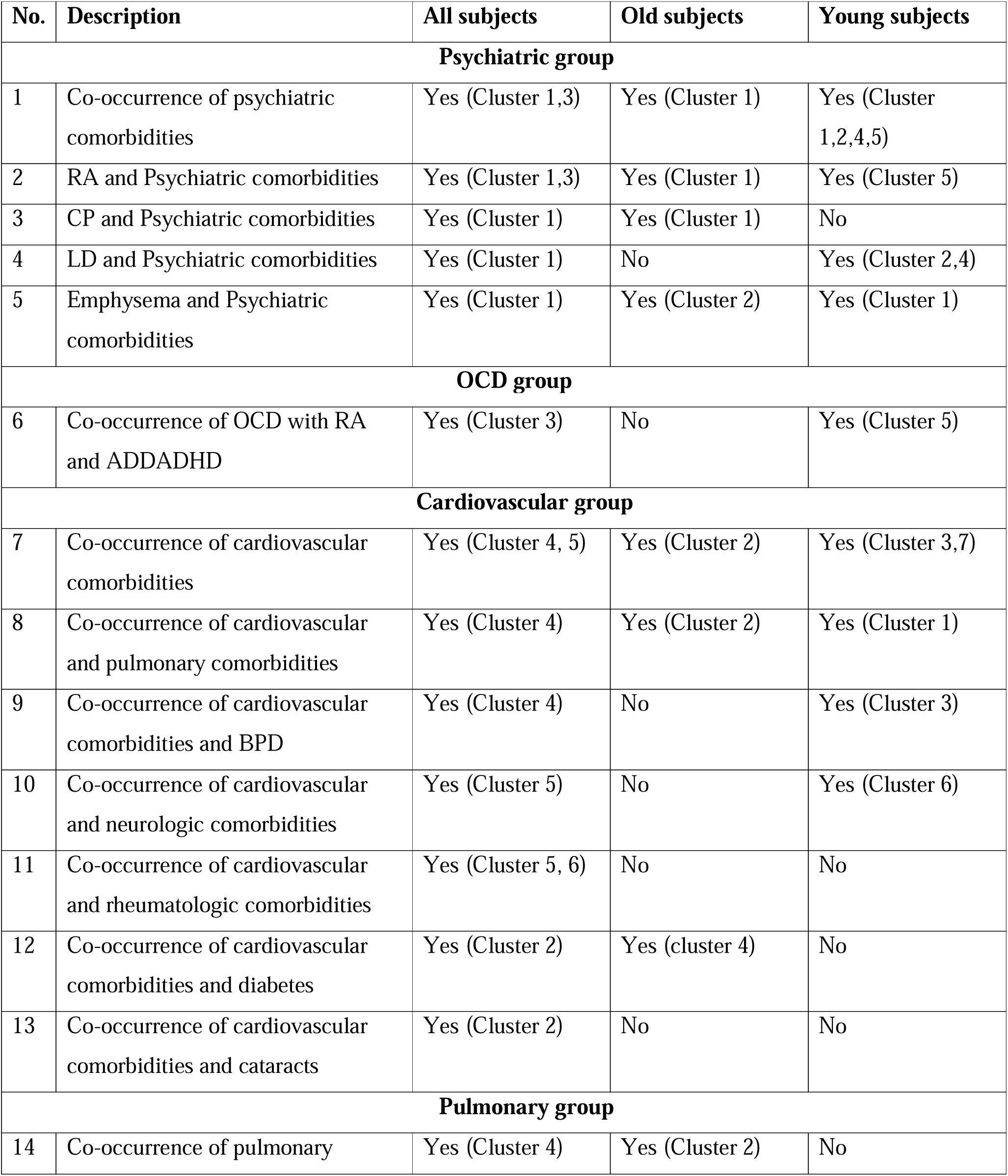
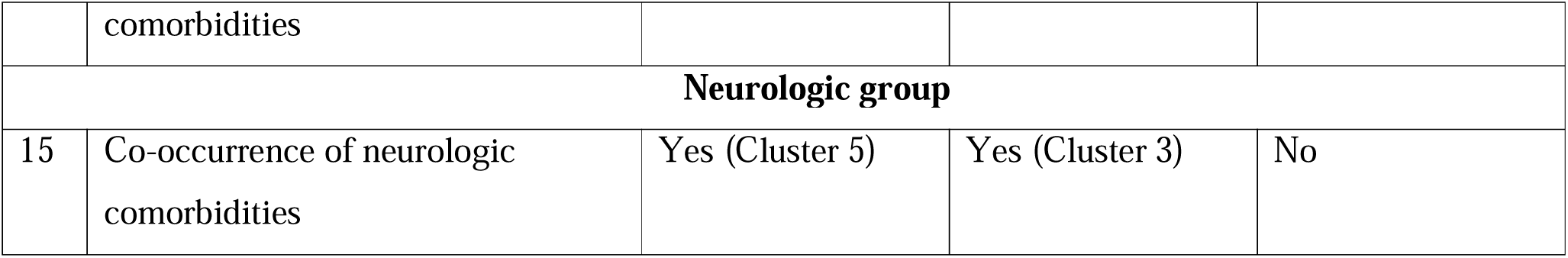
Difference in comorbidity patterns between young and old mTBI subjects.

| No. | Description | All subjects | Old subjects | Young subjects |
| --- | --- | --- | --- | --- |
| **Psychiatric group** |  |  |  |  |
| 1 | Co-occurrence of psychiatric comorbidities | Yes (Cluster 1,3) | Yes (Cluster 1) | Yes (Cluster 1,2,4,5) |
| 2 | RA and Psychiatric comorbidities | Yes (Cluster 1,3) | Yes (Cluster 1) | Yes (Cluster 5) |
| 3 | CP and Psychiatric comorbidities | Yes (Cluster 1) | Yes (Cluster 1) | No |
| 4 | LD and Psychiatric comorbidities | Yes (Cluster 1) | No | Yes (Cluster 2,4) |
| 5 | Emphysema and Psychiatric comorbidities | Yes (Cluster 1) | Yes (Cluster 2) | Yes (Cluster 1) |
| **OCD group** |  |  |  |  |
| 6 | Co-occurrence of OCD with RA and ADDADHD | Yes (Cluster 3) | No | Yes (Cluster 5) |
| **Cardiovascular group** |  |  |  |  |
| 7 | Co-occurrence of cardiovascular comorbidities | Yes (Cluster 4, 5) | Yes (Cluster 2) | Yes (Cluster 3,7) |
| 8 | Co-occurrence of cardiovascular and pulmonary comorbidities | Yes (Cluster 4) | Yes (Cluster 2) | Yes (Cluster 1) |
| 9 | Co-occurrence of cardiovascular comorbidities and BPD | Yes (Cluster 4) | No | Yes (Cluster 3) |
| 10 | Co-occurrence of cardiovascular and neurologic comorbidities | Yes (Cluster 5) | No | Yes (Cluster 6) |
| 11 | Co-occurrence of cardiovascular and rheumatologic comorbidities | Yes (Cluster 5, 6) | No | No |
| 12 | Co-occurrence of cardiovascular comorbidities and diabetes | Yes (Cluster 2) | Yes (cluster 4) | No |
| 13 | Co-occurrence of cardiovascular comorbidities and cataracts | Yes (Cluster 2) | No | No |
| **Pulmonary group** |  |  |  |  |
| 14 | Co-occurrence of pulmonary comorbidities | Yes (Cluster 4) | Yes (Cluster 2) | No |
| **Neurologic group** |  |  |  |  |
| 15 | Co-occurrence of neurologic<br>comorbidities | Yes (Cluster 5) | Yes (Cluster 3) | No |

## Discussion

In this study, we adopted a novel approach of employing graph theoretical principles (Fotouhi et al. 2018; Khan et al. 2019; Sudhakar and Mehta 2024) to identify comorbidity patterns in mTBI subjects. We analyzed the data about mTBI subjects from the TBIMS national database (Traumatic Brain Injury Model Systems Program 2021) five years after the injury. We aim to identify prominent disease associations leading to an enhanced understanding of disease progression in young and old mTBI subjects. Our network analysis of comorbidities post mTBI has yielded interesting associations between disease comorbidities in both young and old cohorts that would eventually contribute to effectively preventing and managing comorbidities.

Prevalence is an important metric that identifies vital comorbidities in a population and is widely quantified in numerous epidemiological studies (Hammond et al. 2019; Sudhakar et al. 2023). Our previous study (Sudhakar et al. 2023) computed the prevalence of various comorbidities after mTBIs. Metric(s) constructed based on graph networks could offer valuable information regarding the presence of comorbidities that play an essential role in disease transitions. Such metrics could also exhibit decreased correlation with prevalence, highlighting the additional information they provide regarding all comorbidities in the population. As mentioned in the results section, betweenness centrality (Figure 3) exhibits a low correlation with prevalence. For example, BPD and emphysema exhibit low prevalence after mTBIs but are characterized by high values of betweenness centrality. On the other hand, CP and anxiety are highly prevalent after mTBI but exhibit low values of betweenness centrality. This could mean that despite exhibiting a low prevalence rate, a specific comorbidity could play a vital role in controlling disease transitions by being associated either directly or indirectly with other comorbidities in the network.

Our results indicate that psychiatric comorbidities tend to co-occur after mTBI incident(s) (Figure 4,6,8), in line with the fact that such illnesses occur at a higher rate in TBI subjects compared to the general public (Shoumitro et al. 1999; Hibbard et al. 2000; Sudhakar et al. 2023). Our study also suggests that OCD is significantly associated with RA in young subjects (Figure 8). While OCD is a psychiatric condition (Alsheikh and Alsheikh 2021), RA is considered to be a rheumatological and autoimmune condition (Hammond et al. 2019; Sudhakar et al. 2023). Endorsing our observations, previous literature suggests that OCD may encompass an inflammatory origin (Mataix-Cols et al. 2018; Alsheikh and Alsheikh 2021). In a nationwide study of Swedish individuals (Mataix-Cols et al. 2018), OCD was associated with an increased number of observing numerous autoimmune conditions. Although individuals in this study exhibited an increased odds of acquiring RA after OCD, the differences were not statistically significant (Mataix-Cols et al. 2018). Additionally, OCD is known to be associated with elevated levels of inflammatory markers such as interleukin (Rao et al. 2015) and TNF-α (Cappi et al. 2012). Therefore, our results are indicative of other studies in the literature that inflammation may contribute significantly to the pathogenesis of OCD (Alsheikh and Alsheikh 2021). OCD was also associated with ADDADHD (Figure 8), and reports suggest that both conditions exhibit a high degree of association and might involve abnormalities in similar brain networks (Brem et al. 2014).

One other association worth mentioning is the link between BPD and cardiovascular diseases (CHF, MI). BPD in the study exhibited comorbidity with cardiovascular conditions (Figure 4) and more so in young subjects (Figure 8). Previous results in the field have corroborated our results (Rossom et al. 2022; Park et al. 2023). In a recent study that quantified long-term cardiovascular disease risk in patients at a primary care centre, subjects with BPD (severe mental illness) exhibited an increased risk (Rossom et al. 2022). Similar results were also observed in a prospective cohort study involving Korean subjects (Park et al. 2023) where young subjects with BPD exhibited an elevated risk for MI. Although poor lifestyle behaviours (BMI, smoking) are postulated as a possible link between BPD and cardiovascular disease risk (Rossom et al. 2022), such a relationship was not observed in the study of the Korean nationwide cohort (Park et al. 2023).

The co-occurrence of emphysema with psychiatric conditions is another notable observation (Figure 4,6,8). In the study, emphysema was associated with depression in both young and old subjects with mTBI history (Figure 4,6,8). Prior literature suggests psychiatric comorbidities (anxiety and depression) are common in subjects with chronic obstructive pulmonary disease (COPD) (Pumar et al. 2014). Possible risk factors for developing psychiatric conditions after COPD include loneliness due to illness severity, dyspnoea and poor physical health (Pumar et al. 2014). COPD subjects with psychiatric conditions are also known to exhibit poor prognosis (Pumar et al. 2014).

Similar to other reports in the literature (Restrepo et al. 2018), we observed that pneumonia is comorbid with COPD (old subjects) (Figure 4,6). Subjects with COPD are at an increased risk of developing pneumonia due to a variety of reasons, including bronchitis, excessive mucus accumulation, etc (Restrepo et al. 2018). CHF also tends to co-occur with pneumonia in our study (Figure 4,6), similar to a previous study (Zhao et al. 2010), which reported that patients with CHF exhibit an increased risk of developing a multitude of comorbidities, including pneumonia, which worsens with age.

We also observed an increased predisposition of psychiatric comorbidities with LD in our study. Among the psychiatric comorbidities, anxiety and PTSD were associated with LD, an observation seen only in young mTBI subjects (Figure 4, 8). Individuals with LD exhibit an increased risk of developing psychiatric conditions and mood disorders, and possible risk factors include tiredness and change in lifestyle (recreation) post-diagnosis (Yovtcheva et al. 2001; Le Strat et al. 2015). Interestingly, PTSD in the veteran population is accompanied by the occurrence of liver cirrhosis, and those veterans with both conditions exhibit dysbiosis of the gut microbiome compared to those without PTSD (Bajaj et al. 2019). Also, liver problems are one of the reasons for increased mortality among veterans with PTSD (Forehand et al. 2019).

Our study has also highlighted potential links between diseases that typically tend to occur in the elderly population. One such example is the connection between diabetes and cataracts (Figure 4) (Kiziltoprak et al. 2019). It’s well documented that subjects with diabetes mellitus exhibit an increased propensity to develop cataracts, an ophthalmologic condition (Kiziltoprak et al. 2019). Abnormal glucose levels in the aqueous humour that nourishes the lens could lead to the accumulation of sorbitol, which eventually results in the deteriorating quality of the lens (Kiziltoprak et al. 2019). Additionally, our study observed a link between LD and cataracts in old subjects (Figure 4,6), documented in a recent survey involving subjects from the UK biobank (Chen et al. 2024). It is assumed that the metabolic modifications caused by the liver condition could eventually lead to the genesis of cataracts (Chen et al. 2024). Another comorbidity association that occurs as people age is the connection between MD and dementia (Figure 4,6) (Tröster and Abbott 2019).

Finally, our study reinforced recent results in the field that cholesterol could be a risk factor for the pathogenesis of OA (Figure 4) (Büchele et al. 2018), a commonly observed rheumatological condition indicating that OA could exhibit a metabolic origin (Farnaghi et al. 2017).

### Limitations of the study and future work

One of the potential drawbacks of the study is the inability to establish directionality when assessing associations between comorbidities. For example, if there is an association between comorbidities A and B in the study, one won’t be able to establish if A preceded B or B preceded A. Currently, it may not be possible to construct a directed disease comorbidity network using the data in the database. All that one can say is the presence or absence of a link between the two comorbidities in study. However, constructing directed disease comorbidity networks would be challenging as it requires tedious documentation of each patient visit and the comorbidities diagnosed during the visit (Khan et al. 2019; Sudhakar and Mehta 2024). Another limitation of the study is the sample size, as the number of mTBI subjects who participated in the NHANES survey ranged from 223-228. The study could be repeated for a larger sample size and different TBI severity forms as a possible future extension. In addition, including control subjects could help understand how the associations established in the study compare with those of the general population.

Another major limitation is the heterogeneity of injury types in mTBIs (Maas 2016; Maas et al. 2017). We tried to address heterogeneity within the mTBI population by constructing separate networks for young and old TBI subjects. In the future, this work can be extended to different TBI pathoanatomic types to combat heterogeneity, as recommended in a recent study (Saatman et al. 2008). Lastly, the presence of ascertainment bias cannot be ruled out, as comorbidities explored in the study could have exhibited a persistent time course and gone unnoticed before the incidence of mTBI (Sudhakar et al. 2023).

## Conclusion

Our study demonstrates the effectiveness of translating disease prevalence data into a graph-theoretical framework. This approach yielded insights into comorbidity patterns following TBI, aligning with previous research, and presents a highly adaptable methodology. The power lies in representing any measurable population characteristic as nodes in a network, with edges signifying associations between them. This not only deepens our understanding but leverages existing data efficiently. Since TBI subjects experience a plethora of comorbidities over time, the study results can be effectively employed for monitoring comorbidities and establishing effective preventive care. Additionally, the approach could be utilized for machine learning-based identification of high-risk patient cohorts (Sudhakar and Mehta 2024).

## List of abbreviations

HBC, LD, PA, BPD, ADDADHD, MI, CHF, OCD, OA, RA, OHC, CP, SD, MD, PTSD, DA, TBIs, mTBIs, NHANES

## Declarations

### Availability of data and materials

All programming scripts used in the analysis and the secondary data that resulted out of the study will be available on reasonable request.

### Author contributions

**Kaustav Mehta**: Methodology, Software, Validation, Formal Analysis, Data Curation, Visualization, Writing - Review and Editing **Shyam Kumar Sudhakar**: Conceptualization, Methodology, Software, Validation, Formal Analysis, Investigation, Resources, Data Curation, Writing - Original Draft, Writing - Review and Editing, Visualization, Supervision, Project Management, Funding Acquisition

## Acknowledgements

This research article used the Traumatic Brain Injury Model Systems National Database, which is supported by funding from the National Institute on Disability, Independent Living, and Rehabilitation Research. NIDILRR is a Center within the Administration for Community Living (ACL), Department of Health and Human Services (HHS). The contents of this research article do not necessarily represent the policy of NIDILRR, ACL, or HHS, and you should not assume endorsement by the Federal Government.

## Supplementary figure legends

**Figure S1**: Phi-correlation coefficient matrix of different comorbidity pairs for all mTBI subjects

The map represents the phi-correlation coefficient values of all comorbidity pairs in mTBI subjects. The green values represent significant associations while the red ones represent in-significant ones

