## Supplementary figures and images for "Exploring Comorbidity Networks in Mild Traumatic Brain Injury Subjects through Graph Theory: A Traumatic Brain Injury Model Systems Study"

### Supplementary Figure 1

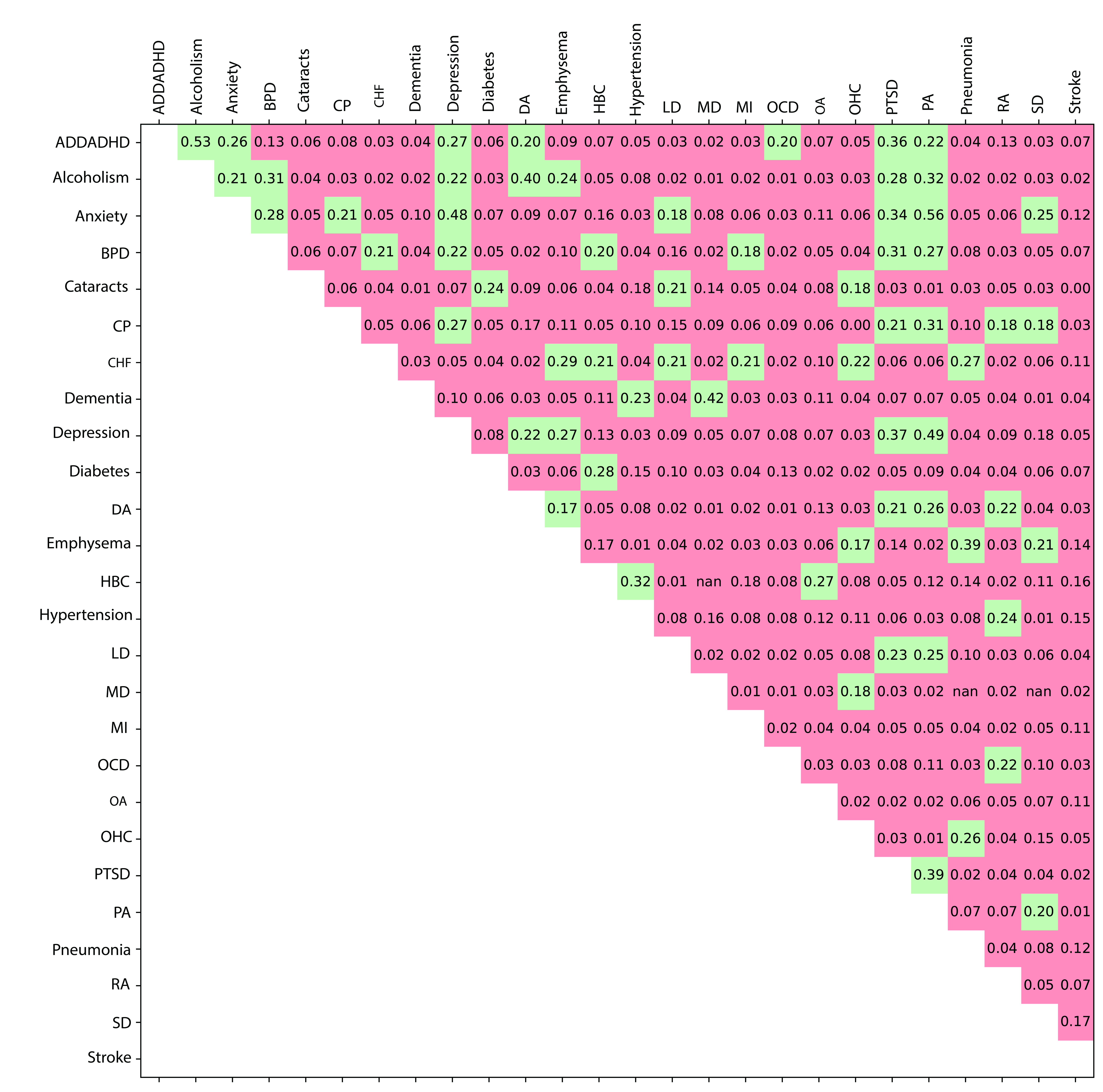
